# An NAMPT Inhibitor Decreases NAMPT Capture by an Antibody Directed against the 5-Phosphoribosyl-1-Pyrophosphate-Binding Loop: A Rational for an NAMPT Occupancy Assay

**DOI:** 10.1101/2024.12.11.628028

**Authors:** Maysoun Shomali, Sandrine Hilairet, Marianne Duhamel, Frederico Nardi, Thomas Bertrand, Magali Mathieu, Rosalia Arrebola, Geraldine Penarier, Samir Jegham, Jerome Arigon, Sylvie Cosnier, Monsif Bouaboula

## Abstract

Nicotinamide phosphoribosyltransferase (NAMPT) catalyzes the rate-limiting step of nicotinamide adenine dinucleotide (NAD) biosynthesis. NAMPT inhibitors (NAMPTi) have been shown to be NAMPT substrates. The resulting NAMPTi-phosphoribose (RP) adduct binds tightly to NAMPT and inhibits the enzyme. Using new NAMPTi, SAR154782, structural analyses performed in this study unveiled a close proximity of the SAR154782-RP complex to the 5-phosphoribosyl-1-pyrophosphate-binding (PRPP-binding) loop within the NAMPT catalytic site. The PRPP-binding loop of NAMPT is subject to conformational flexibility. Interestingly, the PRPP-binding loop domain is oriented to the outer side of the NAMPT dimer and can potentially serve as an antigen-binding site for antibodies. Here we report for the first time that the NAMPTi-RP adduct bound to NAMPT decreases NAMPT capture by an antibody directed against the c-terminal PRPP-binding loop in NAMPT. This finding was then used to explore cellular NAMPT occupancy. The NAMPTi-RP complex displays a sustained, cellular NAMPT occupancy that correlates with the inhibition of NAMPT activity. Moreover, a good correlation between NAMPT occupancy, NAD decrease, and NAMPTi efficacy was observed *in-vivo* in a NCI-H82 small cell lung cancer (SCLC) xenograft model. NAMPT-occupancy assay can be used in clinical settings to better define the optimal dose levels and dose regimen for effective NAMPT inhibition.

## Introduction

There has been a high interest in targeting cancer metabolism pathways such as nicotinamide adenine dinucleotide (NAD) biosynthesis pathways. Nicotinamide phosphoribosyltransferase (NAMPT) is the rate-limiting enzyme for NAD biosynthesis via the salvage route(1). NAMPT transfers a phosphoribosyl group from 5-phosphoribosyl-1-pyrophosphate (PRPP) to NAM to produce nicotinamide mononucleotide (NMN), which is subsequently converted into NAD by nicotinamide mononucleotide adenylyl transferase (NMNAT). Maximal catalytic efficiency of NAMPT requires ATP as a co-substrate. Although ATP is not directly involved in the reaction, phosphoribosyltransferase activity is coupled to concomitant ATP hydrolysis (2, 3).

Because NAMPT is the limiting enzyme for NAD biosynthesis, it represents an attractive target for the development of new cancer therapies. Several classes of NAMPT inhibitors (NAMPTis) have been described (4).

A clear understanding of the NAMPTi profile required for clinical use remains elusive. Despite their potent activities in preclinical models, FK866 and GMX1777 did not show significant clinical activities in cancer patients (5, 6). It remains unclear whether the lack of clinical activity observed with FK866 and GMX1778 was due to a limited exposure precluding NAMPT saturation or a lack of efficacy in treating tumors. Therefore, it is important to develop relevant biomarkers that can assess NAMPT target occupancy and engagement in tumors.

Several compounds are known substrates of NAMPT (7-9). It has been shown through structural studies that an NAMPTi-phosphoribose adduct (NAMPTi-RP) occurs a results of NAMPTi phosphoribosylation, which was detected using recombinant NAMPT. Although it is widely accepted that the adduct is responsible for the inhibition of recombinant NAMPT enzymatic activity, this has not been directly established in a cellular context. Furthermore, the relationships between NAMPT occupancy by the adduct, NAMPT inhibition, and tumor responses remain undefined.

In this study we showed that a novel NAMPTi SAR154782 (10) also serves as an NAMPT substrate. SAR154782 phosphoribosylated adduct (SAR154782-RP) displays a high affinity to NAMPT and is responsible for tight-binding mode NAMPT inhibition. Structural analyses performed in this study unveiled a close proximity of the SAR154782-RP complex to the PRPP-binding loop within the NAMPT catalytic site. The PRPP-binding loop of NAMPT is subject to conformational flexibility (9). Interestingly, the PRPP-binding loop domain is oriented to the outer side of the NAMPT dimer and can potentially serve as an antigen-binding site for antibodies. Taken together, these observations prompted us to investigate whether SAR154782-RP adduct bound to NAMPT interferes with the recognition of NAMPT by an antibody directed against the PRPP-binding loop.

In this study, we showed that SAR154782-RP inhibits the capture of NAMPT by an antibody directed against the PRPP-binding loop of NAMPT. This effect was used to explore the relationships between NAMPT occupancy by SAR154782-RP and in *in-vitro* and *in in-vivo* tumor responses.

## Results

### SAR154782-RP complex unveiled in NAMPT co-crystal structure

SAR154782 is a potent selective NAMPT inhibitor that induce depletion of cellular NAD leading to tumor cell growth inhibition (Fig1A, 1B, 1C, Fig S1A, 1B, 1C).

To gain a better understanding of how NAMPTi such as SAR154782 achieves potent inhibition of NAMPT, we investigated SAR154782 interaction mode of action within NAMPT using co-cristalization studies. The crystal structure of a NAMPT-SAR154782 was solved to 2.1 Å resolution (Fig. 1D upper). This structure shows that SAR154782 binds to the catalytic site within a long, narrow tunnel at the interface of the NAMPT dimer, mostly through hydrophobic interactions.

**Fig. 1.**
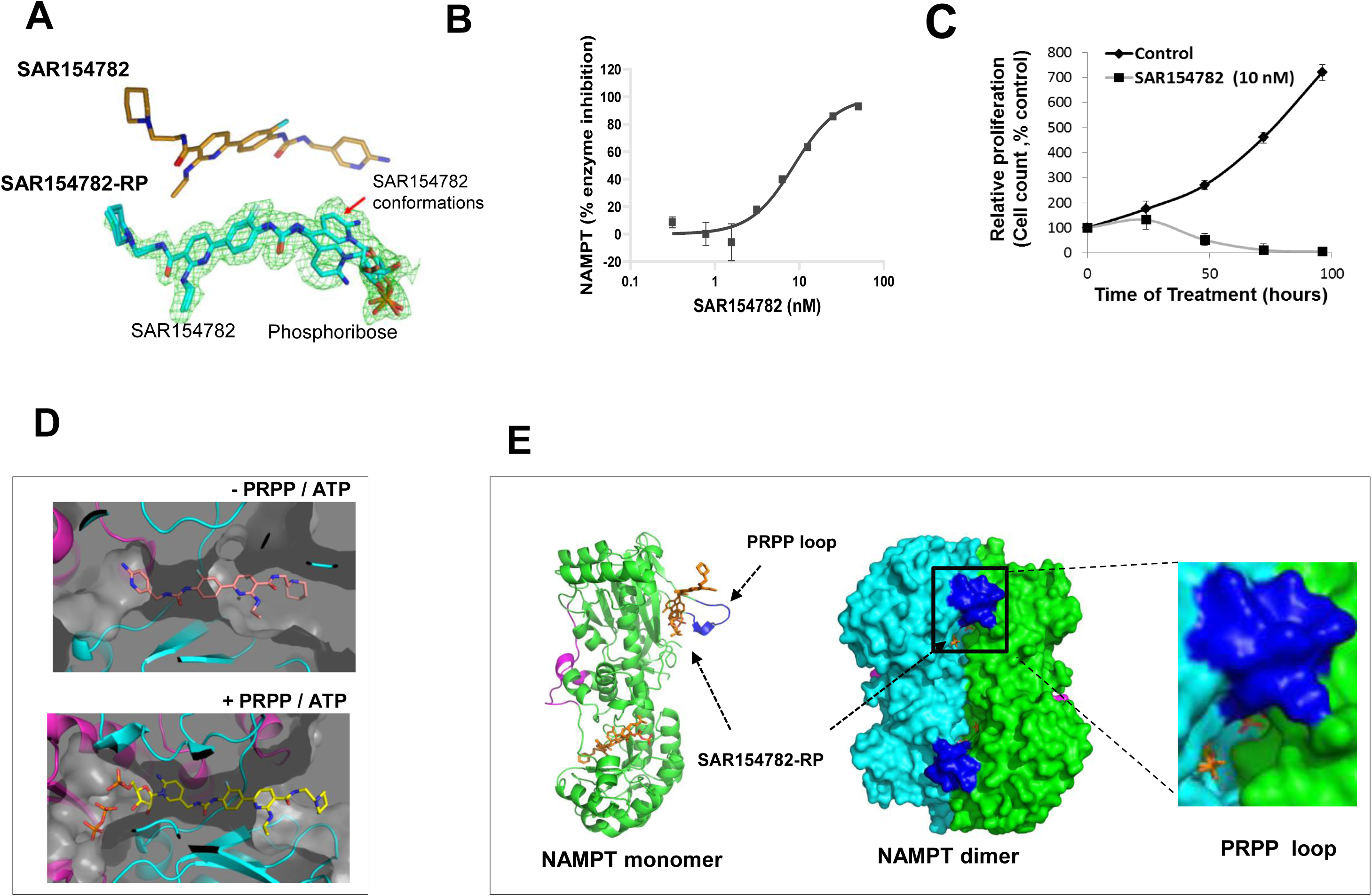
Co-cristal structure of NAMPT with SAR154782 a new NAMPT inhibitor. (A) Chemical structure of SAR54782 (upper panel) and the SAR54782-RP adduct (lower panel). In NAMPT crystal structure, the adduct adopts 2 possible conformations. (B) Dose dependent inhibition of NAMPT enzymatic activity by SAR154782 (NAMPT recombinant protein). (C) Time dependent inhibition of HCT116 cell growth by SAR154782. (D) Co-crystal structure of NAMPT with SAR154782 in absence (upper panel) or presence of PRPP and ATP (lower panel). SAR154782-RP complex can be observed only in presence of PRPP and ATP. (E) Structure of NAMPT dimer. The NAMPT monomers are colored in green and cyan. Two molecules of SAR154782-RP adduct are bound to this structure and are shown in yellow. The structure reveals a close proximity of SAR154782-RP with the PRPP-binding loop. The PRPP-binding loop points toward the opposite side of the NAMPT dimer.

Docking of PRPP in this structure model predicts that SAR154782 pyridine nitrogen and the ribose phosphate of the PRPP substrate are closely apposed, suggesting that the enzyme might be capable of catalyzing the phosphoribosylation of SAR154782. To determine whether SAR154782 is phosphoribosylated in the NAMPT catalytic site, co-crystallization of NAMPT with SAR154782 was carried out under conditions that support the phosphoribosylation reaction. The structure of NAMPT obtained in presence of SAR154782, PRPP, and ATP-Mg^2+^ was determined to 1.88Å (Fig. 1D lower). There is clear density in the channel, corresponding to SAR154782 covalently bound to a ribose-phosphate moiety (SAR154782-RP). The high-resolution data permits to identify two alternate conformations of the inhibitor. Depicted in Fig. 1A, is the Phe193 side-chain which is the more highly populated form. Like SAR154782, SAR154782-RP occupies the nicotinamide side of the tunnel. The ribose-phosphate is located in the PRPP binding pocket and is covalently bound to the nitrogen atom of the pyridine moiety of SAR154782 (Fig. 1B and 1C). Inorganic pyrophosphate (PPi), formed as a reaction product, remains in the binding pocket.

In agreement with the crystal structure findings, an LC/MS-MS analysis of the products formed after a 1 h incubation of NAMPT, PRPP, ATP/Mg^2+^, and SAR154782 demonstrates the formation of a product with a mass of 747, corresponding to the exact mass of the phosphoribosylated SAR154782 adduct identified in the crystal structure (Fig S1D).

### NAMPT inhibition decreases NAMPT capture with an antibody directed against PRPP-binding loop of NAMPT

The co-crystal structure of SAR154782 and NAMPT in the presence of PRPP/ATP revealed a close proximity of the SAR154782-RP complex to the PRPP-binding loop, with a highly hydrophilic zone surrounding the PRPP-binding loop due to the presence of the phosphoribose-SAR154782 linkage (Fig. 1B,1C). It has also been previously shown that the PRPP-binding loop is flexible and subject to conformational changes following binding of the binding NAMPTi-RP adduct (9). Interestingly the NAMPT PRPP-binding loop is oriented towards the outer face of the NAMPT dimer and is susceptible to antibody recognition. Given the close proximity of SAR154782-RP to the PRPP-binding loop and the structural flexibility of PRPP-binding loop, we hypothesized that the SAR154782-RP adduct could interfere with the binding of an antibody directed against the NAMPT PRPP-binding loop.

Initially we started by the identification of an antibody that could recognize the native NAMPT but not the NAMPT preloaded with SAR154782-RP. To preserve the native structure of NAMPT, we performed enzyme-linked immunosorbent assays (ELISAs) with recombinant NAMPT (amino acids [AA] 1–429) serving as the antigen. The NAMPT loaded with SAR154782-RP adduct was generated during the phosphoribosyltransferase step of the NAMPT enzymatic reaction in presence of SAR154782 as a substrate. After completion of the enzymatic reaction in the presence of ethylenediaminetetraacetic acid (EDTA), the enzymatic reaction product was then transferred to an ELISA plate and NAMPT was detected. We evaluated several commercial monoclonal antibodies (mAbs) that were generated against the C-terminus of NAMPT or the entire protein.

Among the panel of antibodies tested, a primary mAb from Adipogen appeared to be highly sensitive to SAR154782-RP adduct for the detection of NAMPT in ELISA (11, 12). As shown in Fig. 2A, a strong ELISA signal corresponding to the capture of recombinant NAMPT by the mAb and detection with a secondary polyclonal Ab was obtained in absence of SAR154782. However, after addition of SAR154782 to the enzymatic reaction, there was a dose-dependent decrease of NAMPT levels detected in ELISA assay. We then investigated whether this decreased signal was related to a decrease of NAMPT recognition and capture by the primary Ab and not due to a limitation with the secondary Ab or the result of non-specific decrease in the ELISA signal. NAMPT enzymatic reaction products were incubated in an ELISA plate pre-coated with the primary mAb. Subsequently, the plate was washed, and the ELISA-well content was harvested using denaturing buffer and analyzed by western blotting using a polyclonal NAMPT Ab. As shown in Fig. 2B, SAR154782 induced a decrease of NAMPT, confirming that SAR154782 blocked recognition and capture of NAMPT by the primary mAb.

**Fig. 2.**
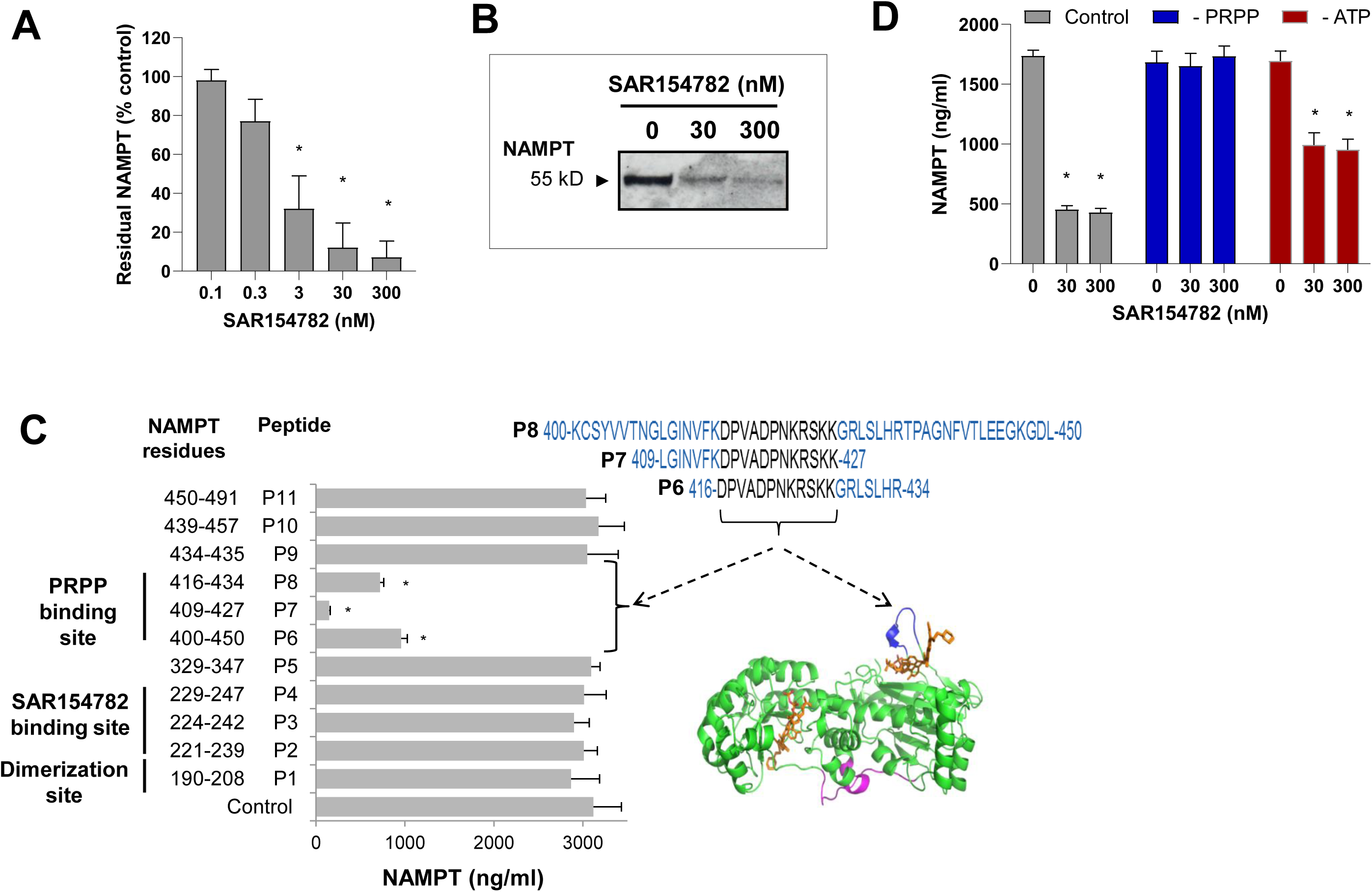
SAR154782 Phosphoribosylation Decreases NAMPT Capture with an Antibody Directed against PRPP-Binding Loop Residues 416–427. (A) SAR154782-RP decreased NAMPT detection in ELISAs. Enzymatic NAMPT reaction products carried out in presence of SAR154782 were transferred to ELISA microplates and NAMPT levels were quantitated. SAR154782 mediated dose-dependent decreases of NAMPT capture. (B) Western blot analysis of NAMPT capture with a primary ELISA mAb. After incubation of enzymatic NAMPT reaction products with a primary ELISA mAb with a subsequent washout step, the ELISA well content was harvested using denaturing buffer, and NAMPT levels were analyzed by western blotting. A strong decrease of the NAMPT was observed following SAR154782 treatment. (C) Epitope mapping of the recognition site by primary ELISA mAb into NAMPT. Competition experiments showing NAMPT-capture inhibition by a primary ELISA mAb in the presence of peptides mapping to different NAMPT domains (left panel). Peptides corresponding to the PRPP-binding loop domain inhibited NAMPT capture (right panel). (D) Effect of SAR154782 phosphoribosylation on NAMPT capture with a primary ELISA mAb. No effect or a low effect of SAR154782 on NAMPT capture in ELISAs when PRPP or ATP was omitted from the enzymatic NAMPT reactions, as indicated. (E) Effect of the SAR154782-RP adduct resident inside the NAMPT enzyme. Removing residual SAR154782 not bound to NAMPT by gel filtration of NAMPT after NAMPT reaction completion shows similar decrease of NAMPT detection in SAR154782 conditions. Each experiment was performed at least twice. Representative data are shown for the mean ± STDEV for a representative experiment. * p<0.05 (compared to untreated control).

We then sought to delineate the NAMPT epitope recognized by the primary antibody, a panel of 11 peptides, mapping to different NAMPT structural domains was used in competition experiments with the primary Ab. The primary mAb was pre-incubated with the different peptides in the presence of full-length NAMPT, followed by extensive washing before adding the secondary Ab. As shown in Fig. 2 C, only the 3 peptides mapping AAs 416-427 of NAMPT were able to inhibit NAMPT capture by this primary mAb, demonstrating that it specifically recognized the PRPP-binding loop (AAs 416-427).

Interestingly, the SAR154782-induced decrease of NAMPT capture was not observed when PRPP was omitted from enzymatic reaction, indicating that this effect was related to the SAR154782-RP adduct, confirming that it interferes with recognition of the NAMPT PRPP-binding loop by the primary ELISA mAb (Fig. 2D). ATP has been shown to contribute indirectly to NAMPTi-RP adduct formation (2). Consistent with an inhibitor role for the adduct, the absence of ATP in the enzymatic NAMPT reaction also affected the SAR154782-induced decrease of NAMPT capture, but to lesser extent (Fig. 2D).

Adding nicotinamide, (the natural substrate of NAMPT) to the enzymatic reaction instead of SAR154782 caused no change in NAMPT capture, indicating that the reaction product NMN did not affect the capture of NAMPT (Not shown). This finding indicated that SAR154782 phosphoribosylation to form the resulting SAR154782-RP adduct specifically inhibited recognition of the PRPP-binding loop by the primary mAb.

An important question is whether the decrease of NAMPT capture observed in ELISAs was due to the adduct locked within the NAMPT enzyme, or to the adduct that could have been released from NAMPT. To address this question, enzymatic NAMPT reaction products were filtered through gel filtration biospin P30 to remove any residual SAR154782 or SAR154782-RP adducts unbound to NAMPT before incubation with the primary ELISA mAb. After filtration, decreased NAMPT capture was still observed (Fig. S2), indicating that this effect was due to adduct bound tightly to NAMPT. Moreover, when SAR154782 was not added during the enzymatic NAMPT reaction, but was added during the ELISA incubation, no effect on NAMPT capture was observed, thereby excluding a role for released SAR154782 (not shown). These results clearly indicated that the SAR154782-RP adduct resident inside NAMPT interfered with the recognition of NAMPT by the mAb directed against the PRPP-binding loop.

### Characterization of SAR154782-RP adduct resident inside NAMPT in HCT116 cells treated with NAMPT inhibitor

Next, we studied whether the cellular NAMPTi-RP adduct bound to NAMPT could also interfere with NAMPT capture in ELISAs. HCT116 cells were pretreated with SAR154782 for 24h, after which the cell extract (lysed under native conditions) was transferred to an ELISA plate for NAMPT quantitation, as described above. As shown in Fig. 3A, SAR154782 treatment dramatically inhibited NAMPT capture in ELISAs. This effect was dose dependent with a maximal effect higher than 80%. However, when a SAR154782-pretreated HCT116 cell extract was assessed for NAMPT expression directly by western blotting under denaturing conditions, no change in NAMPT levels was observed, rulling-out an effect of SAR154782 on the total level of NAMPT protein expression in HCT116 cells (Fig. 3B). As observed with recombinant NAMPT, SAR154782 treatment of HCT116 led to a decrease of native NAMPT capture by the primary ELISA mAb consistent with the interference of the adduct on the binding of mAb directed against the PRPP-binding loop. This decease of NAMPT capture in ELISAs reflects the occupancy of NAMPT by its tightly bound inhibitor, the SAR154782-RP adduct.

**Fig. 3.**
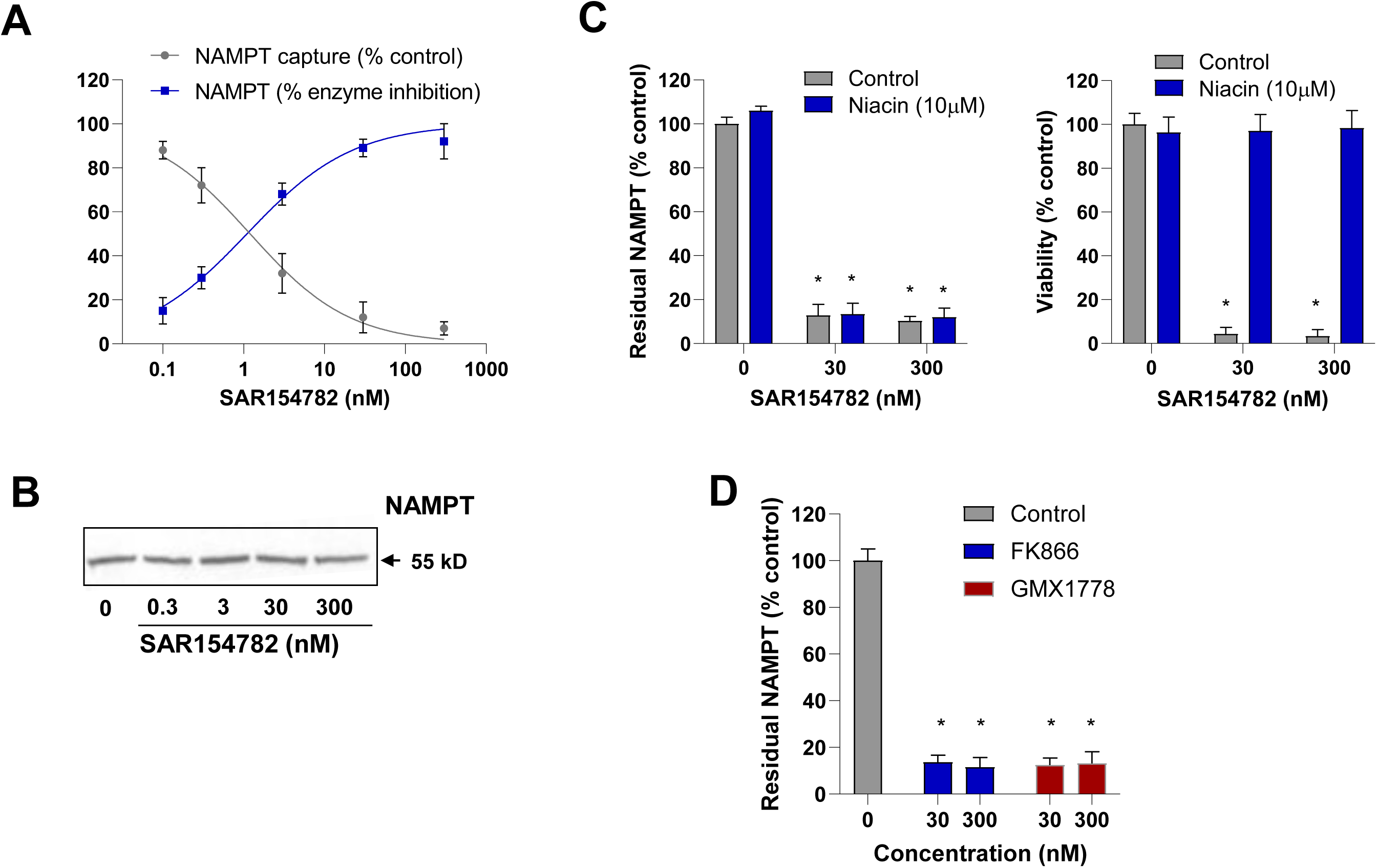
Decrease of NAMPT Capture with a Primary ELISA Ab in HCT116 Cells Treated with NAMPTis. (A) Dose-dependent decrease of NAMPT capture and NAMPT enzymatic activity in cellular extracts from HCT116 cells treated with SAR154782. Parallel to NAMPT capture inhibition results, a dose-dependent inhibition of NAMPT activity by SAR154782 was observed in HCT116 extracts. (B) Western blot analysis of HCT116 cell extracts showed no modulation of NAMPT levels in HCT116 cells treated with SAR154782. (C) Effect of niacin treatment on NAMPT capture (left panel). Treating HCT116 with niacin (10 μM) had no effect on SAR154782 dependent NAMPT-capture inhibition. Niacin reverses the anti-proliferative effect of SAR154782 on HCT116 (right panel). (D) Effects of FK866 and GMX1778 on NAMPT capture. A decrease of NAMPT capture was observed in HCT116 cells treated with FK866 or GMX1778. Representative data are shown for the mean ± STDEV for a representative experiment. * p<0.05, ** p<0.01 (compared to untreated control).

To determine whether NAMPT occupancy by the adduct could inhibit NAMPT activity in treated cells, a cellular extract from SAR154782-treated HCT116 cells was used to measure in parallel, both NAMPT capture in ELISAs and NAMPT enzymatic activity. SAR154782 mediated inhibition of NAMPT capture was dose dependent, with an IC_50_ value in the same range observed for inhibition of enzymatic NAMPT activity (Fig. 3A). The cellular correlation between NAMPT occupancy and inhibition was in agreement with previous findings, confirming the requirement for NAMPT occupancy by the adduct for NAMPT inhibition.

Niacin through another salvage pathway is known to replenish cells with NAD (13) and can reverse the anti-proliferative effects of NAMPTi in HCT116 cells [15-17] (Fig. 3C (right panel)). In HCT116 cells, niacin treatment had no effect on SAR154782-dependent NAMPT capture inhibition in ELISAs, suggesting that neither NAD metabolism, nor NAD-related downstream signaling affected NAMPT occupancy by SAR154782-RP adduct (Fig. 3C left panel).

Next, we studied whether this effect could be observed with NAMPTis other than SAR154782. Interestingly, FK866 and GMX1778, two NAMPT inhibitors with different chemotypes, also showed a strong decrease of NAMPT capture in ELISAs after treatment in HCT116 cells (Fig. 3D). These results suggested that the effect was mediated by adduct formation, given that both GMX1778 and FK866 harbor a 3-pyridyl group that is important for adduct formation (7) and that a GMX1778 adduct has been described previously (7). Our findings are not limited to SAR154782 but may apply to other NAMPT inhibitors.

### The NAMPTi, SAR154782, causes sustained NAMPT occupancy and long lasting inhibition of tumor cell growth

The inhibition of NAMPT capture in ELISAs reflects the occupancy of NAMPT by SAR154782-RP. For simplification for the rest of this study, we will consider NAMPT occupation which is expressed as the percent of NAMPT capture relative to the control (eg 10% capture of cellular NAMPT in Elisa assay will correspond to 90% NAMPT occupancy by its inhibitor). This approach was then used to explore the occupancy of NAMPT by its inhibitor in 2 cell lines H460 and HCT116, which express differing NAMPT levels (Fig. 4A). In both cell lines, the majority of NAMPT was localized in the cytoplasm. Comparing the NAMPT occupancy rates in HCT116 and H460 cells by SAR154782 showed that >80% of NAMPT occupancy could be achieved in both cell lines. However, the EC_50_ for NAMPT occupancy was ∼10-fold lower in HCT116 cells than in H460 cells, consistent with the differences of NAMPT expression observed in these cell lines (Fig. 4B). Thus, higher NAMPT expression was associated with a higher SAR154782 concentration required to achieve the same degree of NAMPT saturation.

**Fig. 4.**
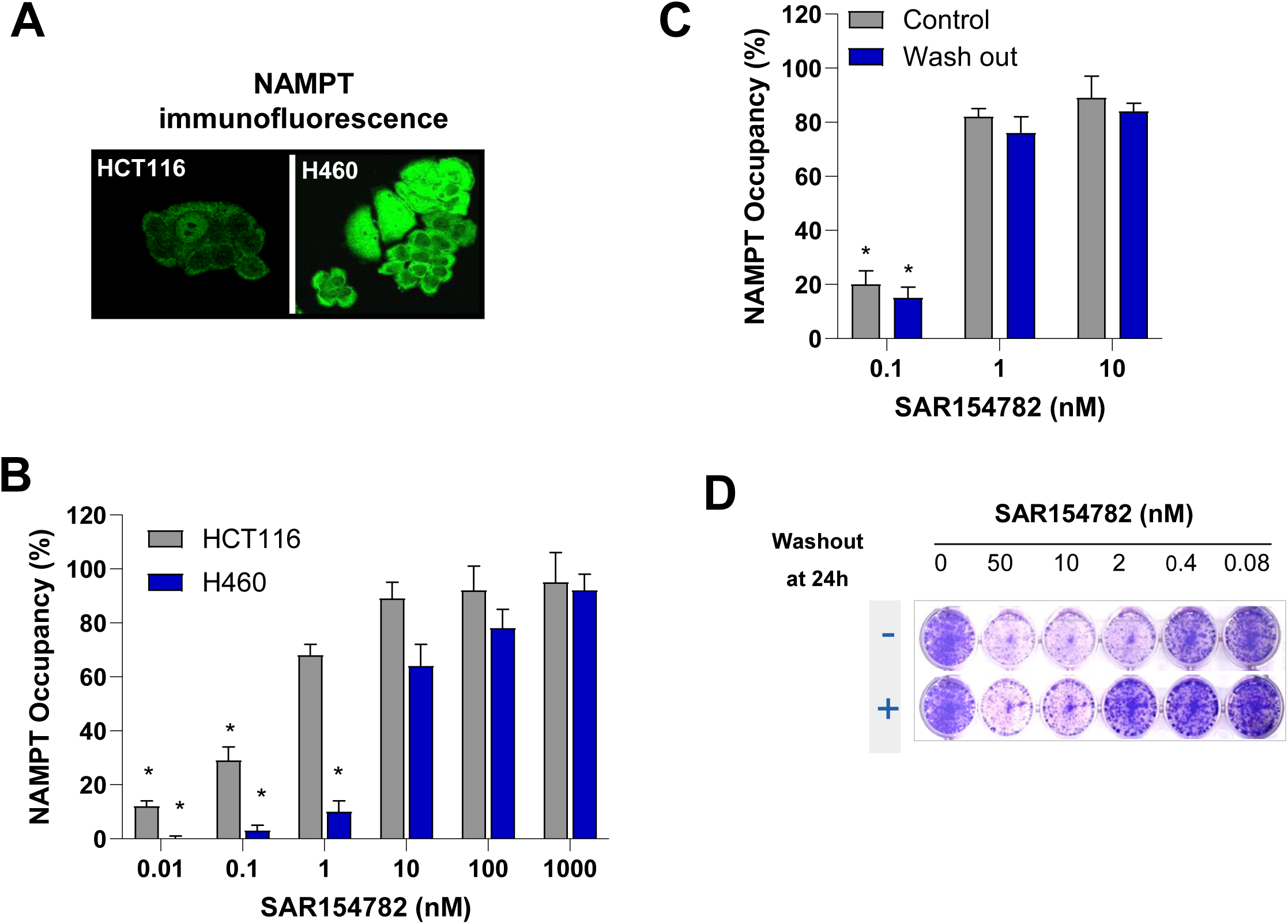
SAR154782 Causes Sustained NAMPT Occupancy and Long-Lasting NAMPT Inhibition in Tumor Cells. (A) Immunofluorescence staining for NAMPT in HCT116 and H460 tumors cells. Higher NAMPT expression levels in H460 tumor cells than in HCT116 tumor cells. (B) In-cellulo NAMPT occupancy by SAR154782-RP. Dose dependent increase of NAMPT occupancy achieving saturation of most cellular NAMPT proteins. Saturation of NAMPT was achieved at lower SAR154782 concentrations in HCT116 cells than in H460 cells, in agreement with the observed differences in NAMPT expression levels in these cell lines. (C) Long-lasting NAMPT occupancy and NAMPT inhibition in HCT116 cells. HCT116 were treated with SAR154782 for 6 h, after which residual SAR154782 was washed out and cells were maintained in culture for an additional 24 h before quantitating NAMPT levels by ELISA. A sustained NAMPT occupancy in HCT116 cells was observed 24 h after an extensive washout of residual SAR154782. (D) Clonal cell growth of HCT116 cells for 7 days after a 6-h treatment and a washout of SAR154782. A slight shift in the SAR154782 concentration required to inhibit HCT116 cell growth after the SAR154782 washout step reflected a long-lasting effect on NAMPT inhibition, consistent with the sustained NAMPT occupancy in HCT116 cells. Each experiment was performed at least twice. The % of NAMPT occupancy is determined according to formula: % Occupancy= 100- (100 * [NAMPT] tretated / [NAMPT] control). Representative data are shown for the mean ± STDEV for a representative experiment.

This approach was also used to explore the reversibility of NAMPT occupancy. HCT116 cells were treated with increasing concentrations of SAR154782 for 6 h and then after a wash-out step, NAMPT was quantitated in ELISAs. A very high NAMPT occupancy rate was still observed 24h later after an extensive wash step was performed to remove unbound SAR154782 (Fig. 4D). These results indicated that in a cellular context, SAR154782 displayed sustained NAMPT occupancy (Fig. 4C). We then explored whether this sustained occupancy could translate in a long-lasting growth inhibition by SAR154782. In clonal-growth assays, HCT116 cells were treated with increasing concentrations of SA154782, washed after a 6h treatment and allowed to grow for an additional 7 days. SAR154782 treatment induced a dramatic inhibition of HCT116 cell growth. A slight shift of the SAR154782 IC_50_ (<5-fold) observed after the washout step was performed most likely reflects a decrease in NAMPT saturation at later time points (after 24 h), since NAMPT occupancy rates were similar up to 24 h, with or without a washout step (Fig. 4D). In agreement with the tight binding, low reversibility mechanism associated with NAMPT inhibition, these data shed light on the role of the phosphoribosylation of the NAMPTi and its long-lasting mode of cellular NAMPT inhibition.

### Relationships between NAMPT occupancy and in *in-vivo* tumor responses

Despite their potent anti-tumor activities in preclinical models, the first-generation NAMPTis FK866 and GMX1777 did not show significant clinical activity. It remains unclear whether these poor clinical outcomes were due to limiting toxicities, sub-maximal NAMPT target occupancy and inhibition, or the absence of anti-tumor efficacies (5, 6).

Building upon SAR154782-induced NAMPT capture inhibition observed in ELISAs, we explored *in vivo* relationships between NAMPT occupancy and tumor responses to SAR154782 in a NCI-H82 small cell lung cancer (SCLC) xenograft model with severe-combined immunodeficiency mice (SCID). NCI-H82-derived tumors were collected after SAR154782 treatment and processed for the quantitation of both NAMPT (ELISA) and NAD (mass spectrometry). SAR154782 induced time- and dose-dependent modulation of NAMPT occupancy (Fig. 5A).

**Fig. 5.**
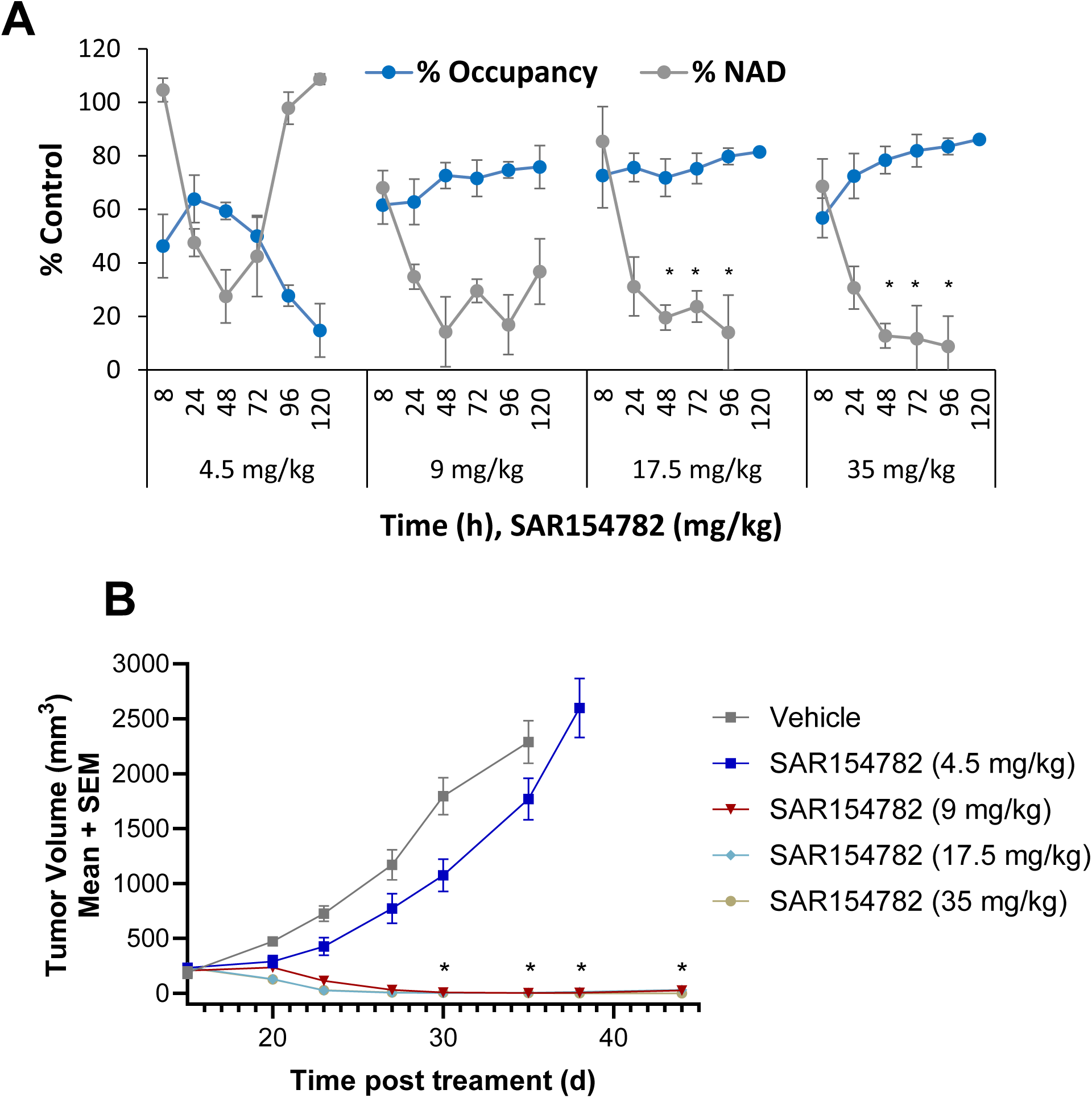
Relationships between NAMPT Occupancy and *In-Vivo* Tumor Responses. (A) Correlation between NAMPT occupancy and NAD-level modulation by SAR154782 in NCI-H82 tumors grafted in SCID mice. After treating mice with increasing doses of SAR154782, tumors were harvested at different time points and processed for the quantitation of both NAMPT and NAD production. Dose- and time-dependent inverse correlations between NAMPT occupancy and NAD modulation were observed. Sustained NAMPT occupancies and NAD decreases were observed for SAR154782 treatments higher than 9 mg/kg. Data are shown for the mean ± STDEV (3 replicates). * p<0.05, * * p<0.01 (compared to untreated control). (B) Anti-tumor efficacy of SAR154782 in SCID mice bearing NCI-H82-derived tumors after a single administration (indicated by arrow). Strong and durable tumor regression was achieved with doses of SAR154782 exceeding 9 mg/kg. Each value represents a median of 6 replicates with standard deviation (STDEV). * p<0.05, * * p<0.01, *** p < 0.001. Complementary analysis of two way anova with repeated measures at fixed level of day with Dunnett’s test versus Vehicle.

In parallel to NAMPT occupancy, we also observed a time- and dose-dependent modulation of NAD levels in these tumors. Indeed, SAR154782 treatment (4.5 mg/kg) induced transient and partial occupancy of NAMPT with a maximal effect of 60% achieved at 24 h post-administration and declined progressively to the baseline at 96 h. Contingent upon NAMPT occupancy, NAD levels decreased in NCI-H82 tumors, showing a ∼75% decrease at 48h post-SAR154782 administration and progressively returned to baseline at 96h post-treatment. At higher doses, SAR154782 caused higher and more sustained NAMPT occupancy over 96h, concomitant with a sustained decrease of NAD. The maximal NAMPT occupancy was ∼80% and corresponded to an NAD decrease of ∼80–90%. In terms of tumor responses, a single SAR154782 administration at 4.5 mg/kg induced delayed tumor growth for at least 5 days, after which the tumor continued growing (Fig. 5B). This partial anti-tumor response was consistent with the partial and transient NAMPT occupancy and NAD decrease. However, at higher doses such as 9, 17.5, or 35 mg/kg, tumor regression was observed within 4–6 days after the onset of SAR154782 administration, in agreement with sustained higher NAMPT occupancy (70–80%) and maximal NAD decrease (80–90%). Remarkably, NCI-H82 tumors continued to regress and were undetectable even 120 days after a single administration of SAR154782 (Fig, 5B, Table 1). These data indicated that in order to achieve tumor regression, it is necessary to have at least a 75–90% NAD decrease and to maintain decreased NAD levels for at least 4–6 days. Accordingly, the inhibition of NAMPT needs to be maintained for at least 4–5 days, consistent with the sustained NAMPT occupancy rate (70–80%) observed at higher doses (Fig. 5B). Based on these data, one would predict that repeated administration of SAR154782 at lower doses such as 4.5 mg/kg, which maintain a sustained NAMPT occupancy (60%) and NAD decrease (75–80%) for 4–6 days, could translate in tumor regression. As expected, when SAR154782 was administered at 4.5 mg/kg in a daily regimen for 5 days, strong tumor regression was achieved (Table 1).

**Table 1.**
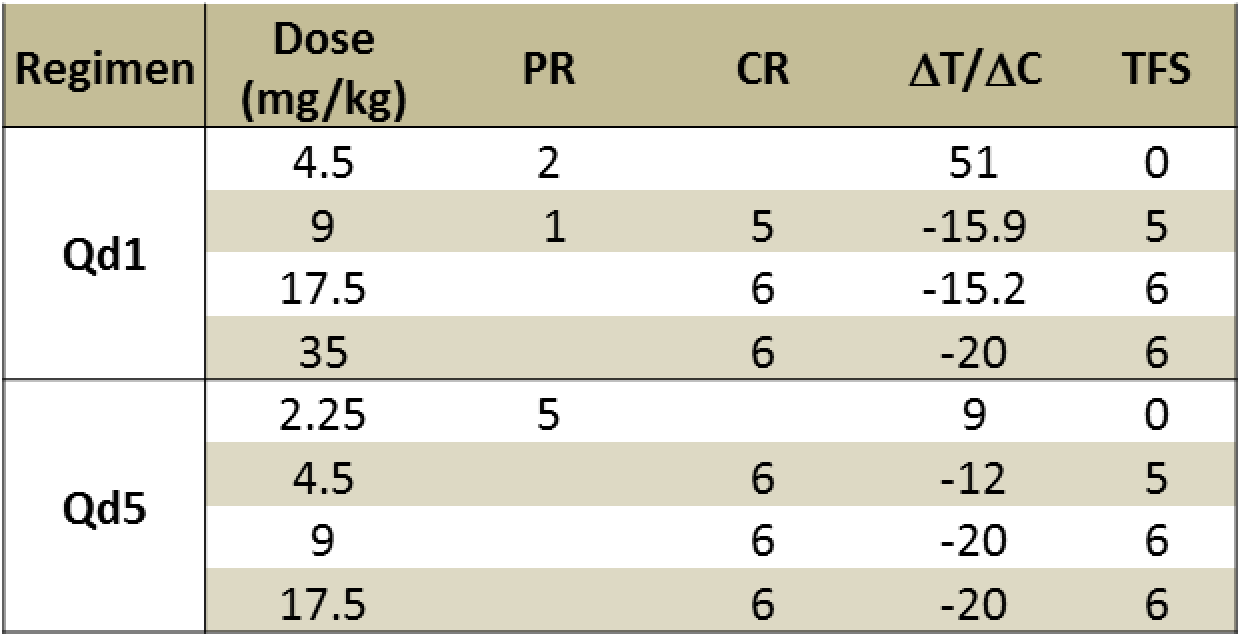
Comparison of the Effects of Single vs. Repeated Dosing Regimens on SAR154782-Dependent Anti-Tumor Efficacy. NCI-H82 tumor-bearing mice were treated with increasing doses of SAR154782 with a daily regimen for 5 days (Qd5) or with a single administration (Qd1). A significant improvement of SAR154782 anti-tumor activities was obtained after repeated dosing with SAR154782 at 4.5 mg/kg. ΔT/ΔC: Ratio of change in tumor volume from baseline median between treated and control groups at day 30 after SAR154782 administration, PR (partial response), CR (complete response), TFS (tumor-free survival: no detectable tumors in mice at 120 days after the first SAR154782 administration)

Taken together, these data validate the correlation between increased NAMPT occupancy, decreased NAD production, and anti-tumor responses. The NAMPT occupancy assay is based on the characteristic ability of the NAMPTi, SAR154782, to form an adduct with phosphoribose that leads to decreased NAMPT recognition by an antibody directed against the PRPP-binding loop (AA416-427). This adduct has a long cellular half-life when bound to NAMPT. In terms of PK/PD relationships, the NAMPT occupancy assay is more directly related to NAMPT inhibition than to tumor NAD decreases, which are time-delayed and may not always reflect the level of NAMPT inhibition particularly given the cross-regulation of metabolic pathways (14). Furthermore we showed that NAMPT occupancy can also be determined in human peripheral blood mononucleare cells (PBMC) treated with SAR154782. Interestingly NAMPT can be detected in human plasma but at lower levels than PBMC. NAMPT occupancy in plasma could also be observed only when PRPP and ATP were added to the SAR154782 treatment, (Fig S3). The NAMPT occupancy assay could be potentially useful in a clinical setting for better defining appropriate NAMPTi plasma concentrations and appropriate regimens required to achieve optimal NAMPT inhibition.

## Discussion

For the first time, we have shown that NAMPT inhibitors such as SAR154782 and FK866 can modulate native NAMPT recognition and capture with an antibody directed against PRPP-binding loop (AAs 416-427). This effect was observed with recombinant NAMPT, as well as with NAMPT in a cellular context. This effect was dependent upon PRPP, confirming the involvement of NAMPT phosphoribosylation. Given the close structural proximity of SAR154782-RP to the NAMPT PRPP-binding loop, these data strongly suggest that the SAR154782-RP complex interfered with binding or recognition of the PRPP-binding loop by the primary mAb. However, it is unclear whether this interference effect was related to steric hindrance, to a conformational change in the PRPP-binding loop, or to the hydrophilic charges of RP. Further investigations are required to delineate this effect.

The decrease of NAMPT capture in ELISAs induced by NAMPTi treatment reflected NAMPT occupancy with SAR154782-RP. In this study, we sought to better understand relationships between NAMPT occupancy rates and tumor responses. NAMPT occupancy with SAR154782-RP was well correlated with the inhibition of NAMPT phosphoribosyltransferase activity.

Identification of biomarkers indicating whether target engagement has been achieved is very critical for clinical trials for new therapeutics. Furthermore, identifying biomarkers for metabolic targets is even more challenging, given the complexities of cross-regulation associated with metabolic pathways. Cellular NAD pools can be regulated by other pathways in parallel to NAMPT activity (14). The modulation of NAD levels is critical for tumor response, but may not fully reflect the degree of NAMPT target inhibition. Hence, NAMPT target occupancy may be a better marker for exploring relationships between NAMPTi pharmacokinetics, NAMPT inhibition, and anti-tumor efficacy. In this study, we validated *in-vitro* and in *in-vivo* decreases of NAMPT capture in ELISAs as NAMPT occupancy assays. We demonstrated that the NCI-H82 SCLC xenograft model, maintaining a NAMPT target inhibition rate of 60–80% for 4–6 days resulted in a 75–90% decrease in NAD production, which was associated with tumor regression. Similar results were obtained in 2 additional in-vivo tumor models involving HCT116 and MX1 cells (data not shown).

It is also important to note that a delay in NAD production inhibition paralleled NAMPT occupation. The delayed NAD decrease was likely related to the NAD metabolization rate in tumor cells. However, this time-kinetic disconnect between NAMPT inhibition and NAD decreases reflects the challenge of using NAD as a direct marker of NAMPT target inhibition. In this study we have shown that NAMPT occupancy could be defined in tumor samples (biopsies from H82 efficacy model) but also in human PBMC or plasma. NAMPT target occupancy testing based on ELISAs could be a very useful approach in a clinical setting to better define appropriate NAMPTi dose and regimens that can translate in optimal NAMPT inhibition.

## Materials and Methods

### Reagents

SAR154782 synthesized as described in (10). FK866 and GMX1778 were obtained from Sigma-Aldrich. HCT116, NCI-H460 and NCI-H82 Cell lines were obtained from ATCC and cultured in DMEM 10% FCS.

### Crystal structure

Crystallization assays were performed on Human NAMPT protein, with N terminal His-tag, was produced in-house in insect cells and purified to 98% purity by chromatography on GSTrap 4B and Superdex 200. The protein was concentrated to 10 mg/mL in 20 mM Hepes, pH 7.5, 100 mM NaCl, 1 mM TCEP and incubated overnight with 1 mM ligand. Crystals are obtained in 100 mM Tris pH8.5, 16% PEG8000. For obtention of NAMPT-SAR154782-RP adduct: The protein was incubated overnight with 1mM of SAR154782, 1mM PRPP, 2mM ATP and 4 mM MgCl_2_. Multiple clustered needle-like crystals rapidly grew and were tested for X-ray diffraction using glycerol as cryoprotectant.

Data were processed using automated methods implemented by GlobalPhasing. The apo-structure of NAMPT, solved by the SGC (PDB code 3IHY) was initially used as a model for molecular replacement. The structures were refined and corrected using Buster (15) and COOT (16).

Final quality checks were done with MolProbity (17). Data collection and data processing statistics can be found in Supporting Information Table S1.

### NAMPT enzymatic assay

For enzymatic activity that were carried out with NAMPT in cell extract, Cell culture of HCT116 cells were treated with increasing concentrations of SAR154782 for 24 hours. For preparation of lysates, 10^7^ cells were resuspended in 100 μl 0.01 mol/l NaHPO4 buffer, pH 7.4, frozen at_-80 °C for 24 h and thawed at room temperature. Cell debris was removed by centrifugation at 23,000g, 20 min at 0 °C. Aliquot of 5 ul of supernatant were incubated in NAMPT enzymatic reaction.

NAMPT enzymatic reactions were carried out in buffer A (50 mM HEPES, pH 7.5, 50 mM NaCl, 5 mM MgCl2) in 96-well microplates. Enzyme solution containing cell extract, 100 μM PRPP, and 1 mM ATP. And 0.5 μCi (1 uM) of [^14^C]-NAM (Amersham, France) was added to the reaction as tracer for NMN biosynthesis. The reaction was incubated at ambient temperature for 120 min before terminated by adding 50 mM EDTA. The produced NMN was determined by [^14^C] NMN precipitation by trichloroacetic acid in presence of BSA carrier, filtration and quantitation of radioactivity in precipitate with microbeta radioactivity counter (wallac system).

For generation of SAR154782-RP adduct, the NAMPT enzymatic reaction using recombinant NAMPT protein was carried out as described earlier but in absence of NAM. The reaction was terminated by adding EDTA and transferred to Elisa plate for quantitation of NAMPT.

### Elisa quantitation of NAMPT

Recombinant NAMPT or NAMPT in cells lysates was quantified using the human intracellular NAMPT ELISA Kit (AdipoGen, Seoul, South Korea), according to manufacturer’s instructions.

Cell treatment and cell extract preparation: HCT116 cells were treated with increasing concentrations of SAR154782 for 24 hours. For preparation of lysates, 10^7^ cells were resuspended in 100 μl 0.01 mol/l NaHPO4 buffer, pH 7.4, frozen at_-80 °C for 24 h and thawed at room temperature. Cell debris was removed by centrifugation at 23,000g, 20 min at 0 °C. Typically cell lysat dilution of 1/5 or 1/20 were incubated in elisa plate to fit with NAMPT elisa standard curve.

For tumors analysis, tumors from treated animals (6 replicates) were pooled in 2 tumors by samples with total of 3 replicates. Tumor lysate were prepared by resuspending tumor fragment 20-100 mg in 500 ul 0.01 mol/l NaHPO4 buffer, pH 7.4. Tumor tissue homogenized with ultra-turrax T18 homogenizer (Sigma Aldrich). The cell debris were removed by centrifugation at 23,000*g*, 20 min, 0 °C. Supernatant dilutions of 1/10 to 1/100 were used in elisa assay.

The % of NAMPT occupancy is determined according to formula: % Occupancy= 100- (100 * [NAMPT] tretated / [NAMPT] control).

### NAMPT peptide mapping

The SAR154782-RP adduct was generated during the NAMPT enzymatic reaction as described earlier. After terminated NAMPT reaction by adding EDTA, 1 μM of synthetic peptides corresponding to different NAMPT domains were added to NAMPT enzymatic reaction and then transferred to elisa wells for quantitation of NAMPT. After incubation with primary capture antibody, the elisa wells were extensively washed and incubated with secondary antibody for detection of NAMPT. The synthetic NAMPT peptides were obtained from thermofisher scientific: Peptide1 (190-LHDFGYRGVSSQETAGIGA-208), Peptide 2 (221-VAGLALIKKYYGTKDPVPG-239), Peptide 3 (224-LALIKKYYGTKDPVPGYSV-242),Pep 4 (229-YYGTKDPVPGYSVPAAEH-247), Pep 5 (329-KKFPVTENSKGYKLLPPY347), Peptide 6 (400KCSYVVTNGLGINVFKDPVADPNKRSKKGRLSLHRTPAGNFVTLEEGKGDL-450),Peptide 7 (409-LGINVFKDPVADPNKRSKK-427), Peptide 8 (416-DPVADPNKRSKKGRL SLHR-434), Pep 9 (434-SLHRTPAGNFVTLEEGKGDL-450), Pep 10 (439-NFVTLEEGKGDLEEYGQDL-457), Peptide 11 (450-LEEYGQDLLHTVFKNGKVTKSYSFDEIRKNAQLNIELEAAHH-491).

### Western blot

HCT116 cell were collected and lysed in RIPA buffer (Pierce) and analyzed for protein expression by western blot using antibodies directed against NAMPT. Monoclonal antibody (OMNI379 Adipogen) was used for detection of total NAMPT in HCT116 cell extract. For the western blot analysis of NAMPT that has been first captured in elisa plate, a polyclonal antibody (Bethyl # A300-779A) was used. Anti-mouse or anti-rabbit secondary antibodies coupled to peroxidase were from Sigma.

### NAMPT gel filtration

The SAR154782-RP adduct was generated during the NAMPT enzymatic reaction using recombinant NAMPT in presence of SAR154782 as described experimental section. In order to purify NAMPT from unbound SAR154782 or unbound SAR154782-RP adduct, NAMPT reaction was terminated by adding 50 mM EDTA, and filtered 3 times through gel filtration biospin P30 (Bio-Rad France). The purified NAMPT was then transferred to NAMPT elisa wells for quantitation as described in experimental section. NAMPT enzymatic activity was also controlled after gel filtration in new enzymatic reaction to confirm the NAMPT inhibition by SAR154782 adduct in comparison with control not previously treatetd with SAR154782.

### Cell growth assay

Cell growth assay was carried out as described in [11]. Briefly, HCT116 cells were seeded into 384-well plates and treated with SAR154782, 24 hours after cell seeding. After a 96-hour incubation period, cells were fixed and stained with nuclear dye. Automated fluorescence microscopy was performed with a GE Healthcare IN Cell Analyzer 1000.

### Immunofluorescence of NAMPT

Analysis of NAMPT expression by confocal fluorescence microscopy. HCT116 an H460 cell lines were cultivated in RPMI 10% SVF in slide dishes. After 48 hours cells were then fixed with 4% paraformaldehyde/ 0.1% saponin. NAMPT was stained with Monoclonal antibody (OMNI379 Adipogen), and FITC coupled anti-mouse secondary antibody (Santa-Cruz). Confocal images were obtained by using a Leica TCS SP2 system with a 63X oil-immersion objective on a Z-stage, and an average of 6 fields with ∼10 cells per field was captured for each group.

### In vivo animal

Tumor cell lines xenografts model: Eight-week-old female severe combined immunodeficiency (SCID) mice were purchased from Charles River. The Committee of Animal Studies at Sanofi approved the protocol for animal experimentation. This protocol and all laboratory procedures complied with French legislation, which implements the European Union directives. NCI-H82 a small cell lung cancer tumor cell line was obtained from DSMZ GmbH. Initially, NCI-H82 cell lines was cultured in RPMI-1640 containing 10% FCS and antibiotics, and implanted subcutaneously in SCID mice (10^7^ cells/mouse). When the tumor reached approximately 1000 mg, it was removed from the donor mouse, cut into fragments (2–3 mm diameter), placed in PBS, and implanted bilaterally with a 12-gauge trocar. Tumor fragments were propagated until stable growth behavior occurred (a stable doubling time), before using in experiments. Distribution was performed using body weight and tumor weight criteria with NewLab Oncology software (NewLab).

For efficacy studies, each model mice bearing tumors of 200 mm3 were treated with dose increase of SAR154782 with an IV route of administration (n=7-8 animals per group). A regimen of single administration or repeated daily administration for five days was used. Drug formulation: SAR154782 IV route: Water/Glucose 5%/ Sodium acetate buffer 50 mM pH4.5/Solutol HS15: 0.25/66.7/4/0.2/28.9. Changes from baseline in tumor volume were used to calculate the median values in treated (ΔT) and control (ΔC) groups. ΔT/ΔC (%) denotes the ratio of medians at any chosen day (the last day before control mice were sacrificed owing to tumor size). ΔT/ΔC values could be translated to activity ratings: ΔT/ΔC < 0: highly active, ΔT/ΔC < 10%: very active, 10% <ΔT/ΔC < 40%: active, ΔT/ΔC > 40%: inactive. When ΔT/ΔC values were negative, the percentage of regression was evaluated. Partial regression (PR) was defined as a decrease > 50% of tumor volume at treatment initiation. Complete regression (CR) was defined as a decrease in tumor volume below the limit of palpation (T = 10 mm^3^). At study end, the number of tumor-free survivors (TFS), corresponding to mice without any detectable tumor, was determined. Both drug-related deaths and maximum percent relative mean net body weight loss were also determined. Median times to reach tumor target size were compared using log-rank or Kruskal–Wallis multiple comparison tests. A body weight loss nadir (mean of group) > 20% or 10% drug-related deaths were considered to indicate an excessive toxic dosage.

Statistical analyses have been performed using internal software Everstat V5 under SAS V8.2 for SUN4. A p-value <0.05 was considered as significant. In order to evaluate the effect of SAR154782 in daily (Qd1) schedule, a two-way ANOVA (Treatment*Day) with repeated measures on day has been performed until day 35 (>50% survival mice in the vehicle group) on the variation of tumoral volume from baseline after rank transformation. Complementary analyses of SAR154782 effect at fixed level of day have been done with Dunnett tests in order to compare treatments to the vehicle group..

### NAD quantification

Tumor Samples from treated mice were collected in Precellys tubes (Bertin) on dry ice and stored at -80°C. Samples were extracted by addition of the internal standard 2-ClAde (2-Chloroadenosine) in the Precellys tube containing the sample, grinding in ACN/H2O 3/1 (v/v), centrifugation, transfer of the supernatant, evaporation to dryness and reconstitution in Ammonium Formate (ForNH4+) 5mM / Acetonitrile (ACN) 95/5 (v/v). Analyses are performed on an automated solid phase extraction system (*Symbiosis, Spark-Holland*) coupled to a LC-MS/MS triple quadrupole mass spectrometer (*TSQ Quantum DiscoMax, Thermo*). The Solid Phase Extraction (SPE) system is used for matrix simplification and sample cleaning. The liquid chromatography (LC) method is based on a C18 reverse-phase column, suitable for polar compounds separation. Column: Atlantis T3 (150 x 2.1 mm, 3 μm / Waters). LC method: gradient from 90/10 H2O/MeOH (ForNH4+) to 30/70 H2O/MeOH (ForNH4+) in 10 min., then return for 10 min. to initial conditions (Flow rate: 0.15 ml/min. and Injection volume: 25 μL). The Mass spectrometric (MS) analysis was performed on a Quantum DiscoMax triple quadrupole (Thermo) with an ESI “ionMax” probe in positive mode. NAD+ and the internal standard 2-ClAde were detected by MS/MS, in multiple reactions monitoring (M.R.M.), by following their specific transitions: NAD+ (664.1 to 136.2). The data were statistically analyzed using the “Everstat 5.0” application.

### Statistical analysis

Statistical analyses have been performed using software Everstat V5 under SAS V8.2 for SUN4. A p-value <0.05 was considered as significant.

## Acknowledgments

The authors would like to thank Claude Bernard for chemical design of SAR154782 as well as Regine Floutard and Isabelle Bentz for technical assistance. The authors also thank Bernard Bourrie for his advises and design of in vivo experiments, as well as Pierre Casellas for initiating the project. The authors acknowledge Alain Jouanen for design of biochemical experiments.

## References

1. Garten A, Petzold S, Korner A, Imai S, Kiess W. Nampt: linking NAD biology, metabolism and cancer. Trends in endocrinology and metabolism: TEM. 2009;20(3):130–8.

2. Burgos ES, Schramm VL. Weak coupling of ATP hydrolysis to the chemical equilibrium of human nicotinamide phosphoribosyltransferase. Biochemistry. 2008;47(42):11086–96.

3. Burgos ES, Ho MC, Almo SC, Schramm VL. A phosphoenzyme mimic, overlapping catalytic sites and reaction coordinate motion for human NAMPT. Proceedings of the National Academy of Sciences of the United States of America. 2009;106(33):13748–53.

4. Galli U, Travelli C, Massarotti A, Fakhfouri G, Rahimian R, Tron GC, et al. Medicinal chemistry of nicotinamide phosphoribosyltransferase (NAMPT) inhibitors. Journal of medicinal chemistry. 2013;56(16):6279–96.

5. von Heideman A, Berglund A, Larsson R, Nygren P. Safety and efficacy of NAD depleting cancer drugs: results of a phase I clinical trial of CHS 828 and overview of published data. Cancer chemotherapy and pharmacology. 2010;65(6):1165–72.

6. Holen K, Saltz LB, Hollywood E, Burk K, Hanauske AR. The pharmacokinetics, toxicities, and biologic effects of FK866, a nicotinamide adenine dinucleotide biosynthesis inhibitor. Investigational new drugs. 2008;26(1):45–51.

7. Watson M, Roulston A, Belec L, Billot X, Marcellus R, Bedard D, et al. The small molecule GMX1778 is a potent inhibitor of NAD+ biosynthesis: strategy for enhanced therapy in nicotinic acid phosphoribosyltransferase 1-deficient tumors. Molecular and cellular biology. 2009;29(21):5872–88.

8. Zheng X, Bauer P, Baumeister T, Buckmelter AJ, Caligiuri M, Clodfelter KH, et al. Structure-based discovery of novel amide-containing nicotinamide phosphoribosyltransferase (nampt) inhibitors. Journal of medicinal chemistry. 2013;56(16):6413–33.

9. Oh A, Ho YC, Zak M, Liu Y, Chen X, Yuen PW, et al. Structural and biochemical analyses of the catalysis and potency impact of inhibitor phosphoribosylation by human nicotinamide phosphoribosyltransferase. Chembiochem: a European journal of chemical biology. 2014;15(8):1121–30.

10. Bernhart C, Bouaboula M, Casellas P, Jegham S, Arigon J, Combet R, et al. Nicotinamide derivatives, preparation thereof and therapeutic use thereof. Google Patents; 2010.

11. Kover K, Tong PY, Watkins D, Clements M, Stehno-Bittel L, Novikova L, et al. Expression and regulation of nampt in human islets. PloS one. 2013;8(3):e58767.

12. Laiguillon MC, Houard X, Bougault C, Gosset M, Nourissat G, Sautet A, et al. Expression and function of visfatin (Nampt), an adipokine-enzyme involved in inflammatory pathways of osteoarthritis. Arthritis research & therapy. 2014;16(1):R38.

13. Kirkland JB. Niacin status, NAD distribution and ADP-ribose metabolism. Current pharmaceutical design. 2009;15(1):3–11.

14. Opitz CA, Heiland I. Dynamics of NAD-metabolism: everything but constant. Biochemical Society transactions. 2015;43(6):1127–32.

15. Bricogne G. Direct phase determination by entropy maximization and likelihood ranking: status report and perspectives. Acta crystallographica Section D, Biological crystallography. 1993;49(Pt 1):37–60.

16. Emsley P, Cowtan K. Coot: model-building tools for molecular graphics. Acta crystallographica Section D, Biological crystallography. 2004;60(Pt 12 Pt 1):2126-32.

17. Davis IW, Leaver-Fay A, Chen VB, Block JN, Kapral GJ, Wang X, et al. MolProbity: all-atom contacts and structure validation for proteins and nucleic acids. Nucleic acids research. 2007;35(Web Server issue):W375-83.

