## Supplemental figures for "An NAMPT Inhibitor Decreases NAMPT Capture by an Antibody Directed against the 5-Phosphoribosyl-1-Pyrophosphate-Binding Loop: A Rational for an NAMPT Occupancy Assay"

### Slide 1
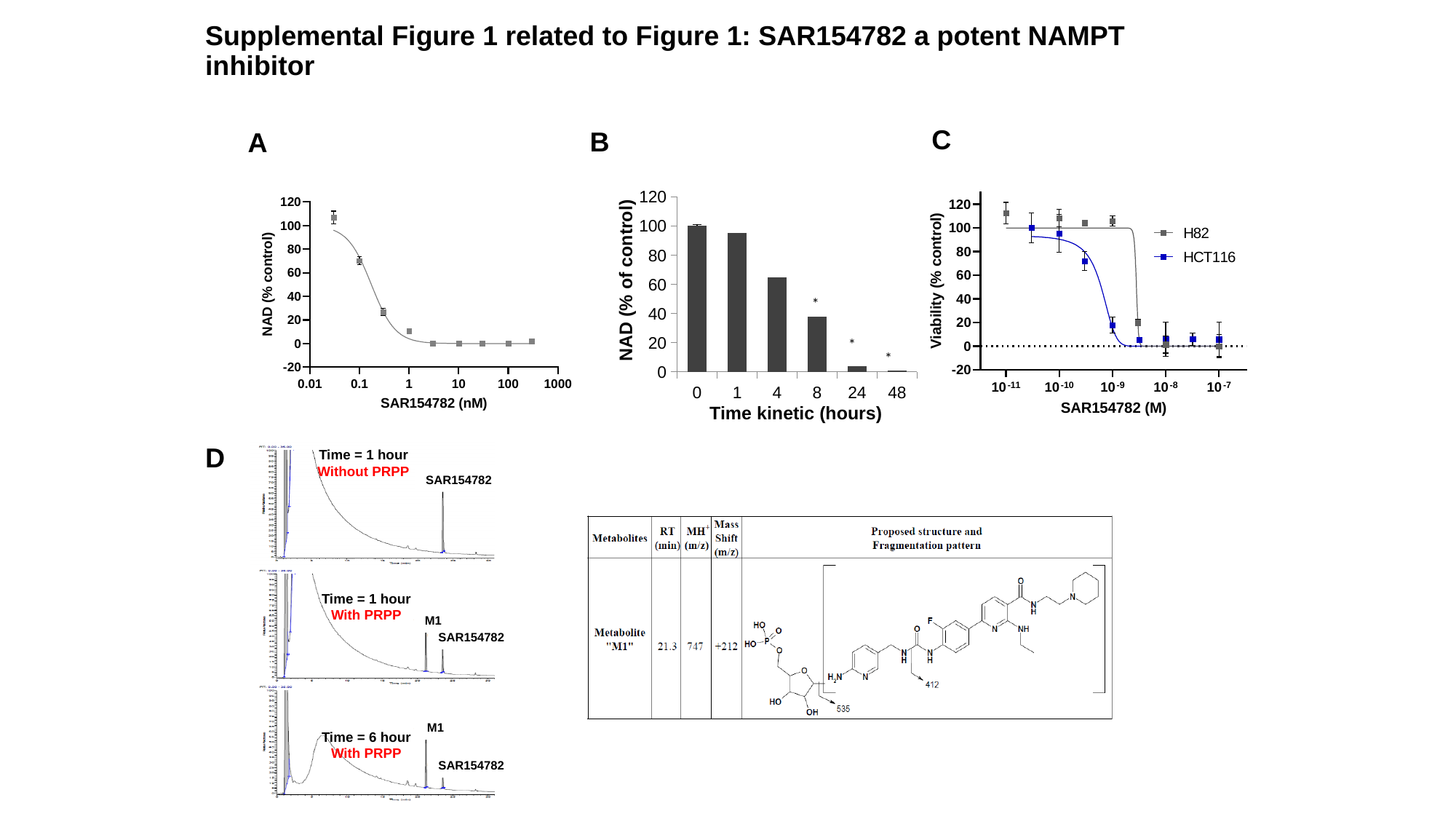

#### Supplemental Figure 1 related to Figure 1: SAR154782 a potent NAMPT inhibitor
C
B
A
###### Chart
| Category | |
|---|---|
| 0 | 100.0 |
| 1 | 95.0 |
| 4 | 65.0 |
| 8 | 38.0 |
| 24 | 4.0 |
| 48 | 1.0 |NAD (% of control)
*
*
*
Time kinetic (hours)
1
D
Time = 1 hour
Without PRPP
SAR154782
Time = 1 hour
With PRPP
M1
SAR154782
M1
Time = 6 hour
With PRPP
SAR154782

### Slide 2
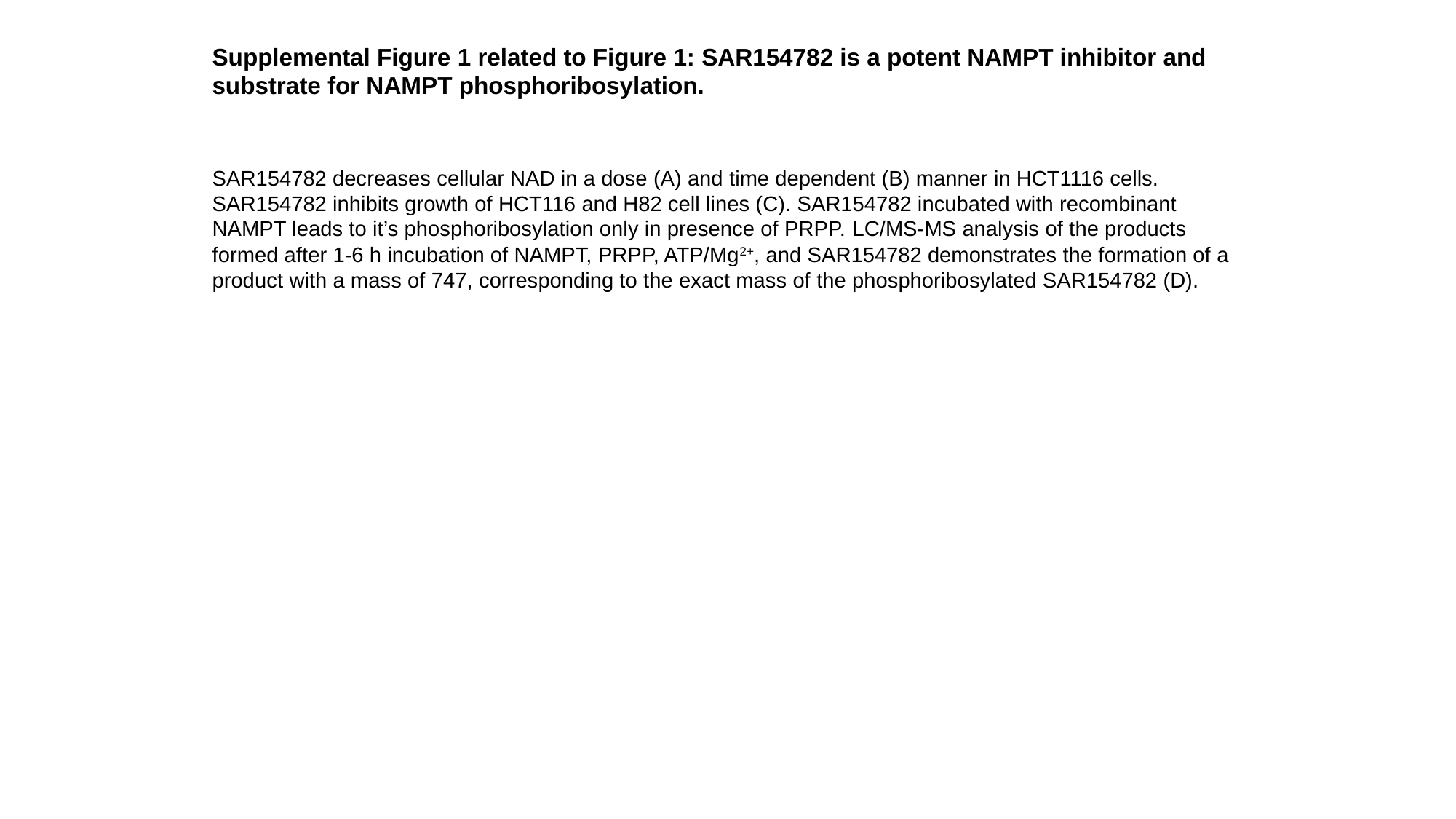

Supplemental Figure 1 related to Figure 1: SAR154782 is a potent NAMPT inhibitor and substrate for NAMPT phosphoribosylation.
SAR154782 decreases cellular NAD in a dose (A) and time dependent (B) manner in HCT1116 cells. SAR154782 inhibits growth of HCT116 and H82 cell lines (C). SAR154782 incubated with recombinant NAMPT leads to it’s phosphoribosylation only in presence of PRPP. LC/MS-MS analysis of the products formed after 1-6 h incubation of NAMPT, PRPP, ATP/Mg^2+^, and SAR154782 demonstrates the formation of a product with a mass of 747, corresponding to the exact mass of the phosphoribosylated SAR154782 (D).

### Slide 3
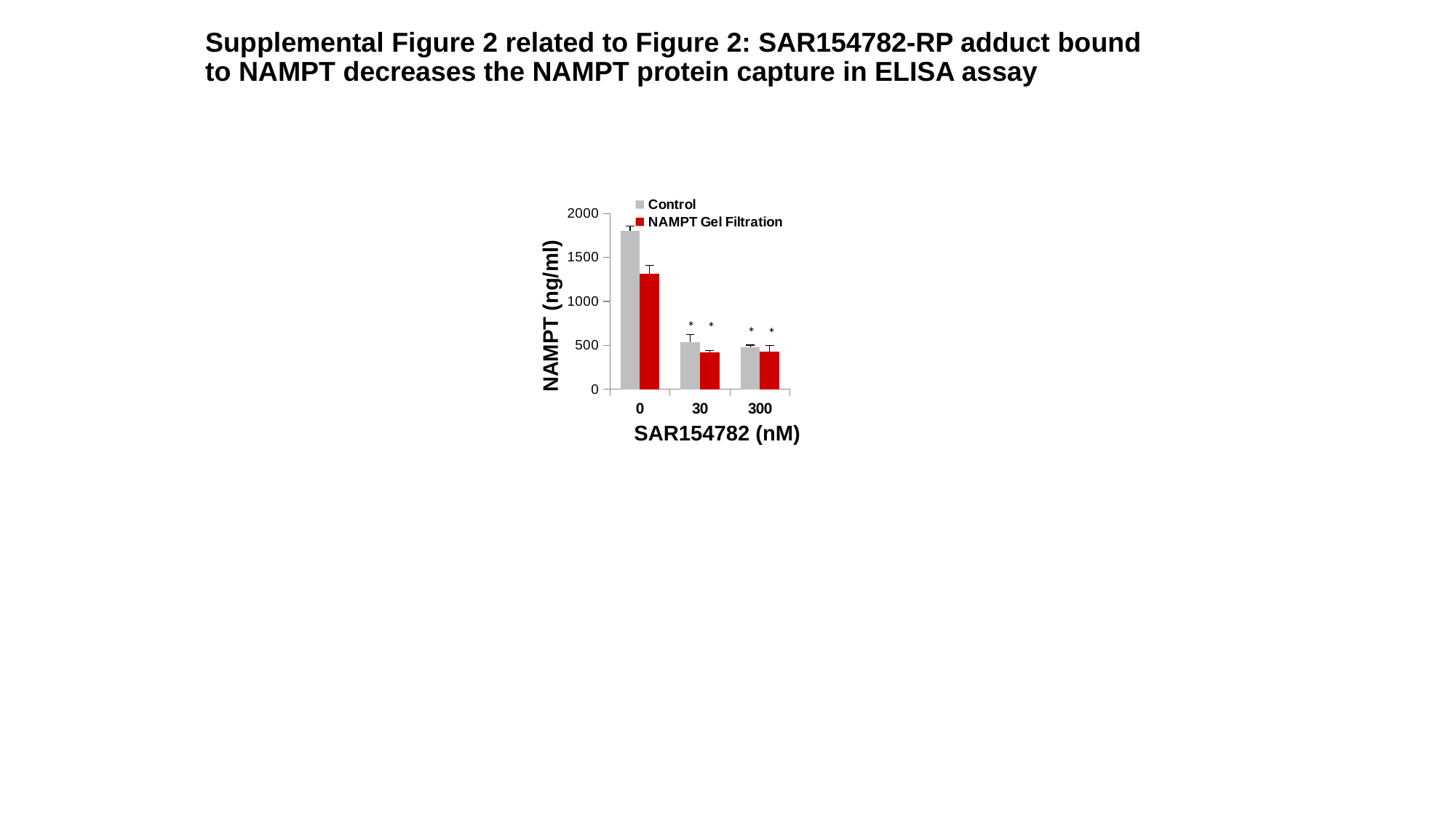

#### Supplemental Figure 2 related to Figure 2: SAR154782-RP adduct bound to NAMPT decreases the NAMPT protein capture in ELISA assay
###### Chart
| Category | Control | NAMPT Gel Filtration |
|---|---|---|
| 0 | 1800.0 | 1310.0 |
| 30 | 540.0 | 420.0 |
| 300 | 480.0 | 430.0 |NAMPT (ng/ml)
*
*
SAR154782 (nM)
*
*

### Slide 4
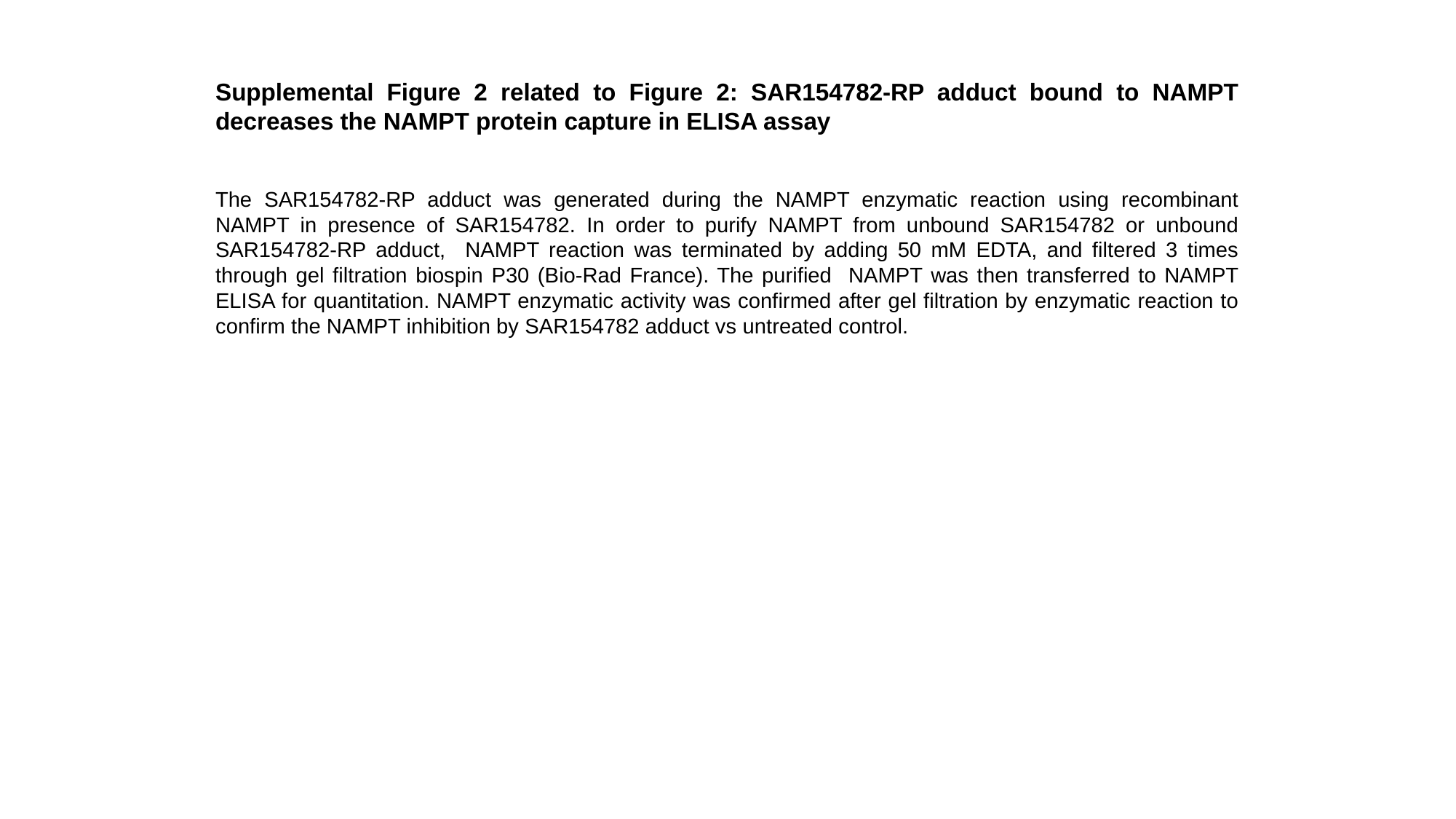

Supplemental Figure 2 related to Figure 2: SAR154782-RP adduct bound to NAMPT decreases the NAMPT protein capture in ELISA assay
The SAR154782-RP adduct was generated during the NAMPT enzymatic reaction using recombinant NAMPT in presence of SAR154782. In order to purify NAMPT from unbound SAR154782 or unbound SAR154782-RP adduct, NAMPT reaction was terminated by adding 50 mM EDTA, and filtered 3 times through gel filtration biospin P30 (Bio-Rad France). The purified NAMPT was then transferred to NAMPT ELISA for quantitation. NAMPT enzymatic activity was confirmed after gel filtration by enzymatic reaction to confirm the NAMPT inhibition by SAR154782 adduct vs untreated control.

### Slide 5
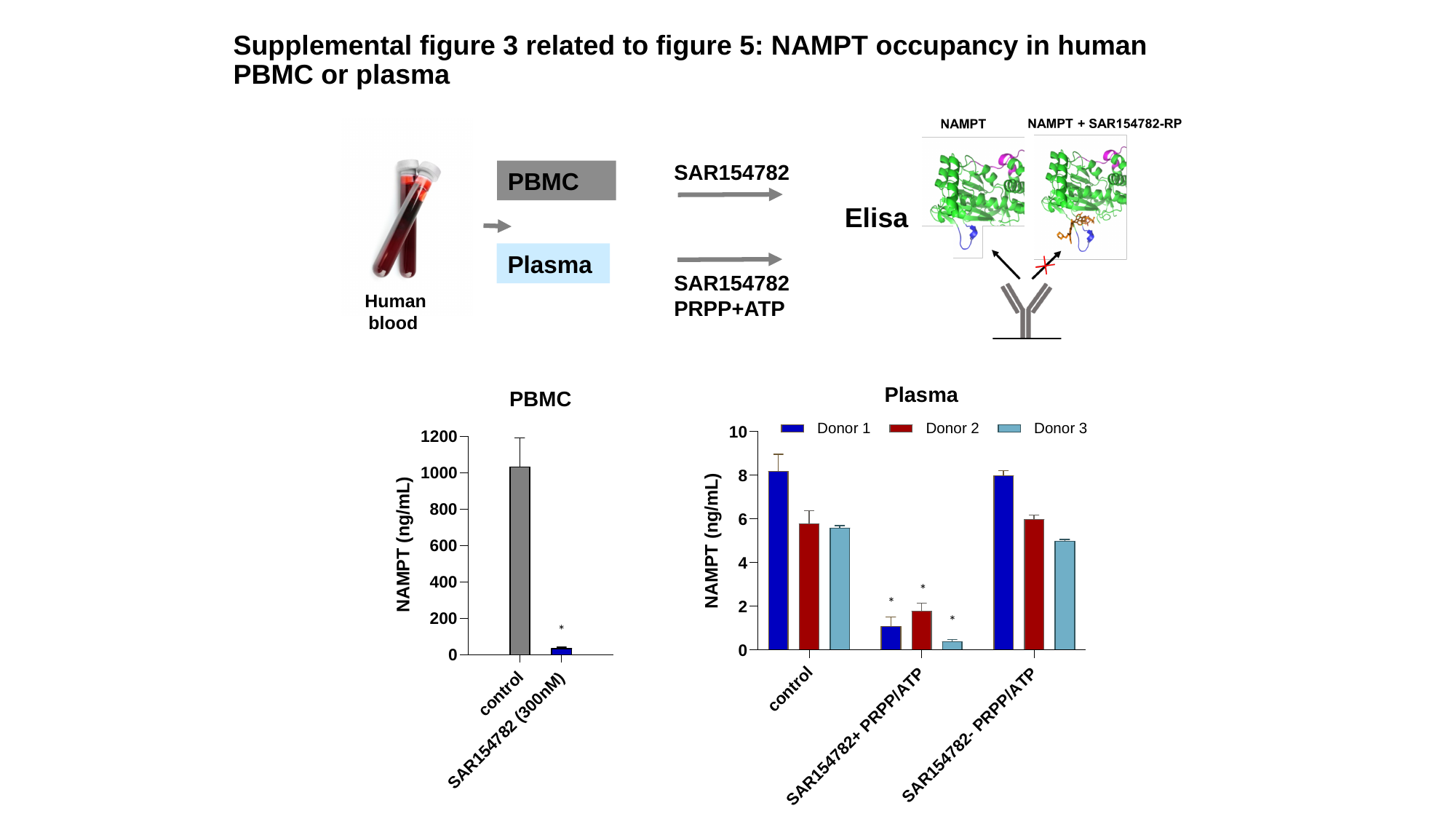

#### Supplemental figure 3 related to figure 5: NAMPT occupancy in human PBMC or plasma
SAR154782
PBMC
Elisa
Plasma
SAR154782
PRPP+ATP
Human blood
*
*
*
*

### Slide 6
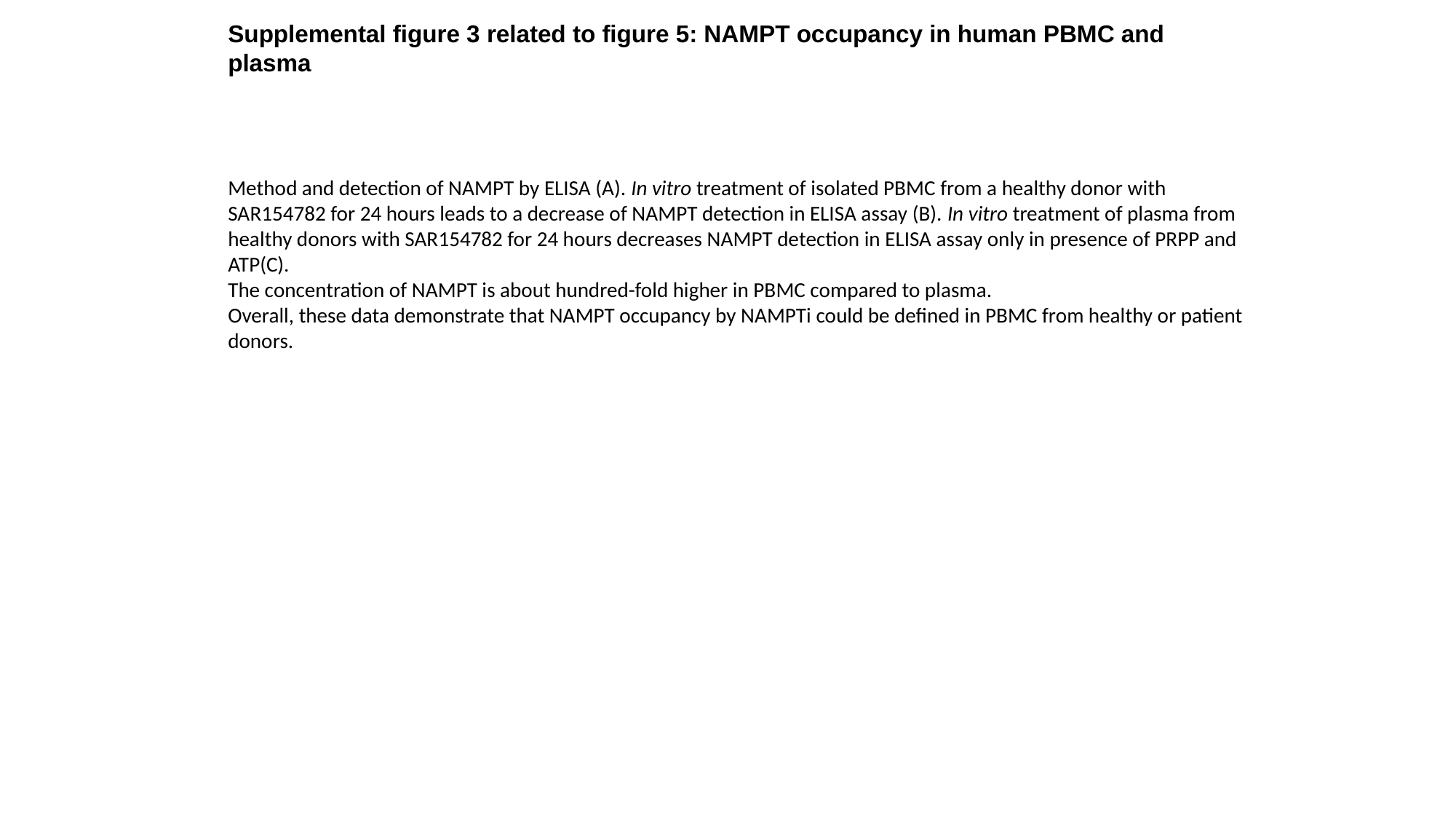

Supplemental figure 3 related to figure 5: NAMPT occupancy in human PBMC and plasma
Method and detection of NAMPT by ELISA (A). In vitro treatment of isolated PBMC from a healthy donor with SAR154782 for 24 hours leads to a decrease of NAMPT detection in ELISA assay (B). In vitro treatment of plasma from healthy donors with SAR154782 for 24 hours decreases NAMPT detection in ELISA assay only in presence of PRPP and ATP(C).
The concentration of NAMPT is about hundred-fold higher in PBMC compared to plasma.
Overall, these data demonstrate that NAMPT occupancy by NAMPTi could be defined in PBMC from healthy or patient donors.
